# VolcaNoseR – a web app for creating, exploring, labeling and sharing volcano plots

**DOI:** 10.1101/2020.05.07.082263

**Authors:** Joachim Goedhart, Martijn S. Luijsterburg

**Affiliations:** Swammerdam Institute for Life Sciences, Section of Molecular Cytology, van Leeuwenhoek Centre for Advanced Microscopy, University of Amsterdam, P.O. Box 94215, NL-1090 GE Amsterdam, The Netherlands. Email: | Twitter: @joachimgoedhart; Department of Human Genetics, Leiden University Medical Center, Einthovenweg 20, 2333 ZC, Leiden, The Netherlands. Email: | Twitter: @luijsterburglab

## Abstract

Comparative genome- and proteome-wide screens yield large amounts of data. To efficiently present such datasets and to simplify the identification of hits, the results are often presented in a type of scatterplot known as a volcano plot, which shows a measure of effect size versus a measure of significance. The data points with the largest effect size and a statistical significance beyond a user-defined threshold are considered as hits. Such hits are usually annotated in the plot by a label with their name. Volcano plots can represent ten thousands of data points, of which typically only a handful is annotated. The information of data that is not annotated is hardly or not accessible. To simplify access to the data and enable its re-use, we have developed an open source and online web tool with R/shiny. The web app is named VolcaNoseR and it can be used to create, explore, label and share volcano plots (https://huygens.science.uva.nl/VolcaNoseR). When the data is stored in an online data repository, the web app can retrieve that data together with user-defined settings to generate a customized, interactive volcano plot. Users can interact with the data, adjust the plot and share their modified plot together with the underlying data. Therefore, VolcaNoseR increases the transparency and re-use of large comparative genome- and proteome-wide datasets.

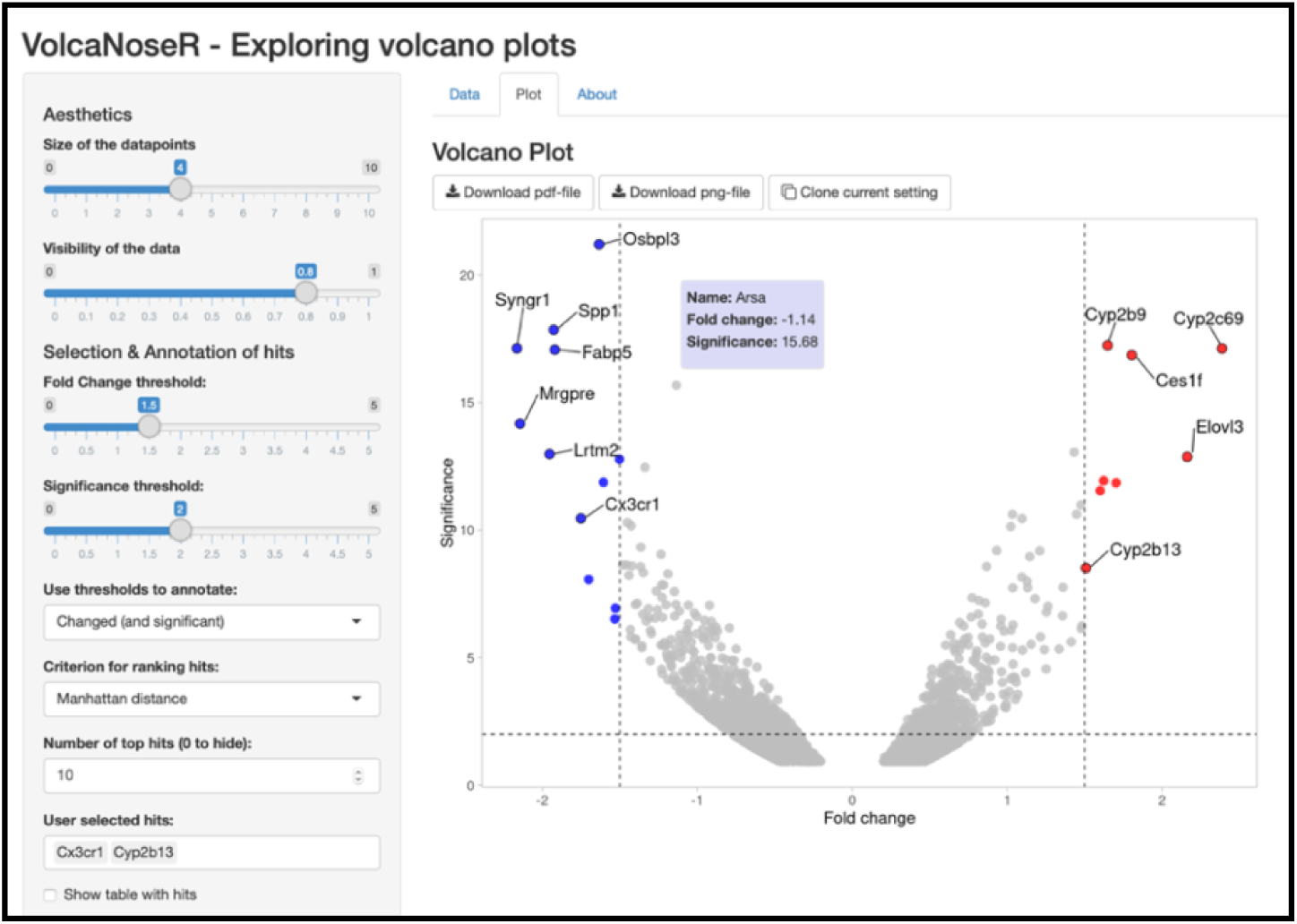

## Introduction

The volcano plot visualizes complex datasets generated by genomic screening or proteomic approaches. It is essentially a scatter plot, in which the coordinates of datapoints are defined by effect size and statistical significance (Li, 2012; Cui and Churchill, 2003). Volcano plots typically show the data of hundreds to ten thousands of genes or proteins. Examples of such datasets are gene expression changes measured by RNA-seq (Becares et al., 2019), genome-wide loss-of-function CRISPR screens (Drainas et al., 2020), or mapping the interactome of proteins-of-interest by mass spectrometry (van der Weegen et al., 2020). Although volcano plots are based on rich datasets, only a handful of data points are usually labeled with a gene or protein name. This enables the visual identification of hits and simplifies the interpretation of the complex dataset. Nevertheless, the data points that are not annotated may be of equal interest. Therefore, it is highly desirable to have easy access to the information of all data points from such large datasets.

Volcano plots are typically generated using commercial software or with software that requires coding skills. A viable alternative is provided by dedicated free web apps that allow users to generate plots through a graphical user interface (GUI). Several web apps are available (Singh et al., 2016; Naumov et al., 2017), but these do not generate interactive plots and only have limited options for customization and annotation. Therefore, we decided to generate a web-based online tool for generating volcano plots, similar to previously plotting apps (Goedhart, 2020; Postma and Goedhart, 2019). Here, we report an open source web app for generating, exploring, labeling and sharing Volcano plots. The web app is created with R/Shiny and is dubbed VolcaNoseR. Below we discuss the features of the app.

### Availability, code & issue reporting

The VolcaNoseR webtool is available at: https://huygens.science.uva.nl/VolcaNoseR or at (as long as the bandwidth limit is not reached): http://goedhart.shinyapps.io/VolcaNoseR/

The code was written in R using R (https://www.r-project.org) and Rstudio (https://www.rstudio.com). To run the app several freely available packages are required: shiny, ggplot2, magrittr, dplyr, ggrepel, shinycssloaders, DT and RCurl. The code of version 1.0.2 reported in this manuscript is archived at Zenodo.org: https://doi.org/10.5281/zenodo.3855569

Up-to-date code and new release will be made available on Github, together with information on running the app locally: https://github.com/JoachimGoedhart/VolcaNoseR

The Github page of VolcaNoseR is the preferred way to communicate issues and request features (https://github.com/JoachimGoedhart/VolcaNoseR/issues). Alternatively, the users can contact the developers by email or <u>Twitter</u>. Contact information is found on the “About” page of the app.

### Data input and format

The data can be supplied via upload of a text file with different delimiters, including the Comma Separate Values (CSV) format. Upload of excel workbooks with multiple sheets is also supported. Alternatively, a CSV file from an online data repository can be used through a URL. To demonstrate the features of the app, example data is included of which the details can be found elsewhere (Becares et al., 2019; Gillingham et al., 2019).

After data upload, the user selects the columns that hold the information on the fold change (for the x-coordinate) and the significance (for the y-coordinate). Selecting a column with gene or protein names is optional.

### Data visualization

A typical volcano plot shows the log_2_ of the fold change on the x-axis and minus log_10_ of the p-value on the y-axis. The data is shown as dots and their size and transparency can be adjusted. The position of the individual points is defined by these coordinates. By hovering over the data points the information about the data can be accessed immediately and dynamically. When the pointer (mouse) is near a data point, the x- and y-coordinate and the name is retrieved, providing the user with easy access to the underlying data. In some cases, it may be desirable to display a 90 degrees rotated volcano plot. This option is available and will depict the fold change on the y-axis and the significance on the x-axis.

### Thresholds and hits

The user can set threshold values for the fold change and the significance. The threshold values are indicated by dashed lines in the plot and used to classify the data as ‘unchanged’, ‘decreased’ or ‘increased’. The data are colored according to this classification and this can be shown in a legend. The ‘top hits’ can be automatically detected and ranked based on a number of criteria. The default criterion is the Manhattan distance (|ΔX|+|ΔY|) of the data from the origin (0,0). The other criteria are Euclidean distance (SQRT(ΔX^2^+ΔY^2^)), absolute fold change or significance. The data are sorted based on the selected criterion and the 10 top-ranking data points are selected. The number of top ranking data points can be adjusted by the user.

It is possible to annotate only ‘increased’ or ‘decreased’ or all significantly changed (‘increased’ and ‘decreased’) data points. The top-ranking hits are shown in the plot and there is an option to list them in a table. Finally, the user can manually search and select the names of genes or proteins of interest, which will be annotated in the plot and added to the table.

The standard colors to indicate ‘unchanged’, ‘increased’ and ‘decreased’ are respectively grey, red and blue. Another color combination that is available is grey, blue and green. Users can also define their own color scheme.

### Output

Users can customize the titles for the axes and their size. The plot that is generated by the app can be directly retrieved by drag-and-drop from the web browser. In addition, the plot can be downloaded as a PNG or PDF file. The PNG is a lossless bitmap format. The PDF allows for downstream processing/editing with software that can handle vector-based graphics.

### Sharing data and plot settings

All settings that are defined in the user interface can be stored as a URL, as was previously implemented for PlotsOfData and PlotTwist (Postma and Goedhart, 2019; Goedhart, 2020). When the data is retrieved from an external online resource, this hyperlink is included in the URL. The URL with settings is sufficient to (i) launch the app, (ii) retrieve the data, and (iii) plot the data according to user-defined settings. Once the plot is available, it can be adjusted and a new URL reflecting the new settings can be obtained. This feature enables transparent reporting of all the data and simplifies re-use of the data. We illustrate this feature with data from proteomic screens that we have recently published (van der Weegen et al., 2020). These data are deposited and publicly available at the data repository zenodo.org, doi: <u>10.5281/zenodo.3713174</u>. Volcano plots are generated with VolcaNoseR using the data from the CSV files in the repository. Next, the URL that encodes all necessary information was generated using the ‘clone current setting’ button. With this unique URL, the data is retrieved and a plot is generated by VolcaNoseR based on the parameters that are stored in the URL. For instance, this URL produces an interactive plot of which a static version is shown in figure 2A:

https://huygens.science.uva.nl:/VolcaNoseR/?data=5;;Difference_CSB_GFP;p_value_CSB_GFP;Gene_names&vis=4;0.8;1.5;2;increased&can=10;;;&layout=;;;;;log2;minus_log10;X;600;800&label=TRUE;GFP-CSB-WT vs GFP-NLS;TRUE;Enrichment (log2);Significance (-log10 p- value);;24;24;18;6;&url=https://zenodo.org/record/3713174/files/CSV_1-GFP-CSB-WT_vs_GFP-NLS.csv

**Figure 1:**
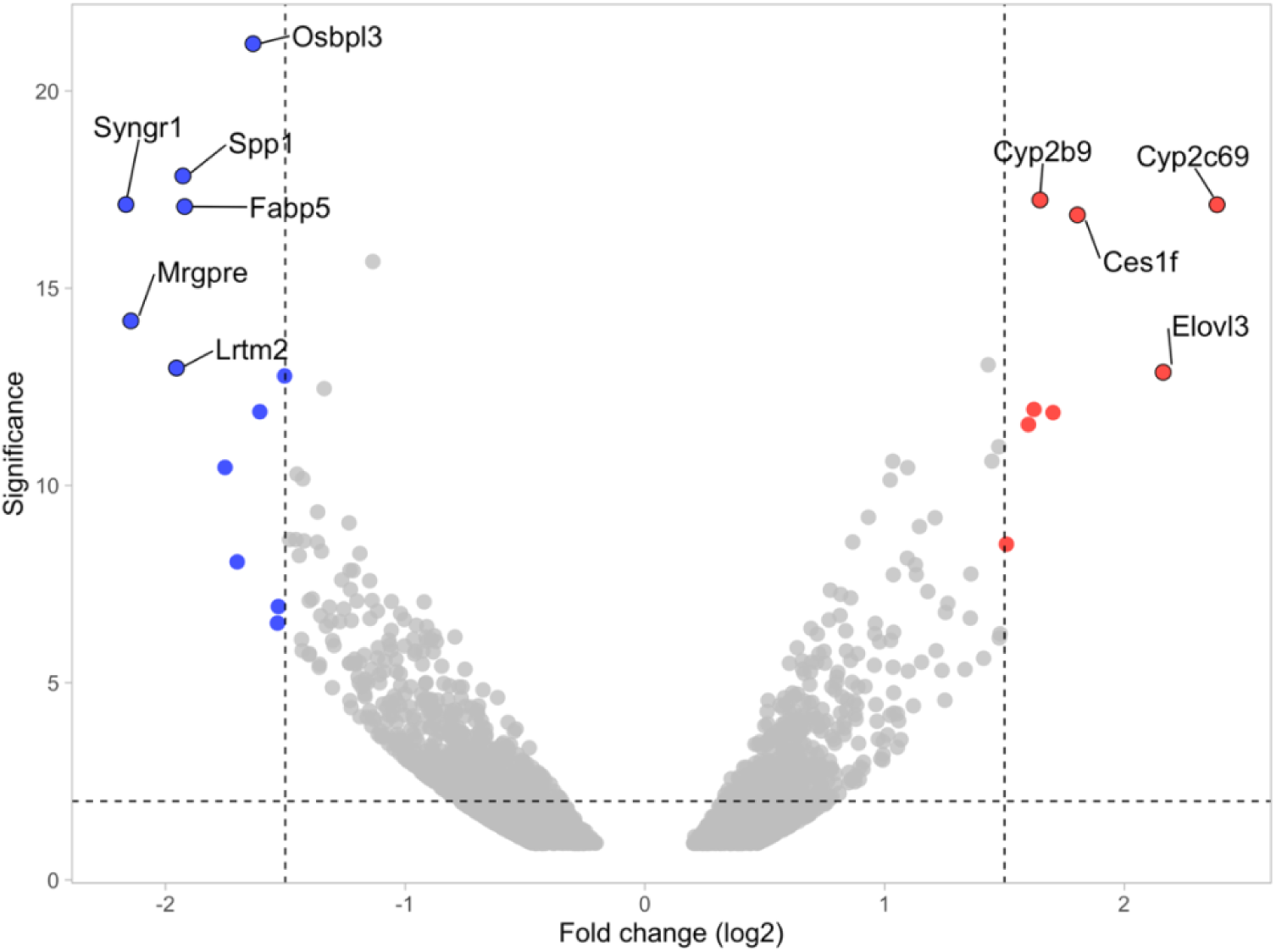
The standard output of the VolcaNoseR app for example dataset 1. The annotated dots are the ten datapoints that have the largest (Manhattan) distance from the origin and are above the thresholds indicated by the dashed line. Direct access to an interactive plot and all the data is provided by <u>this hyperlink</u>.

**Figure 2:**
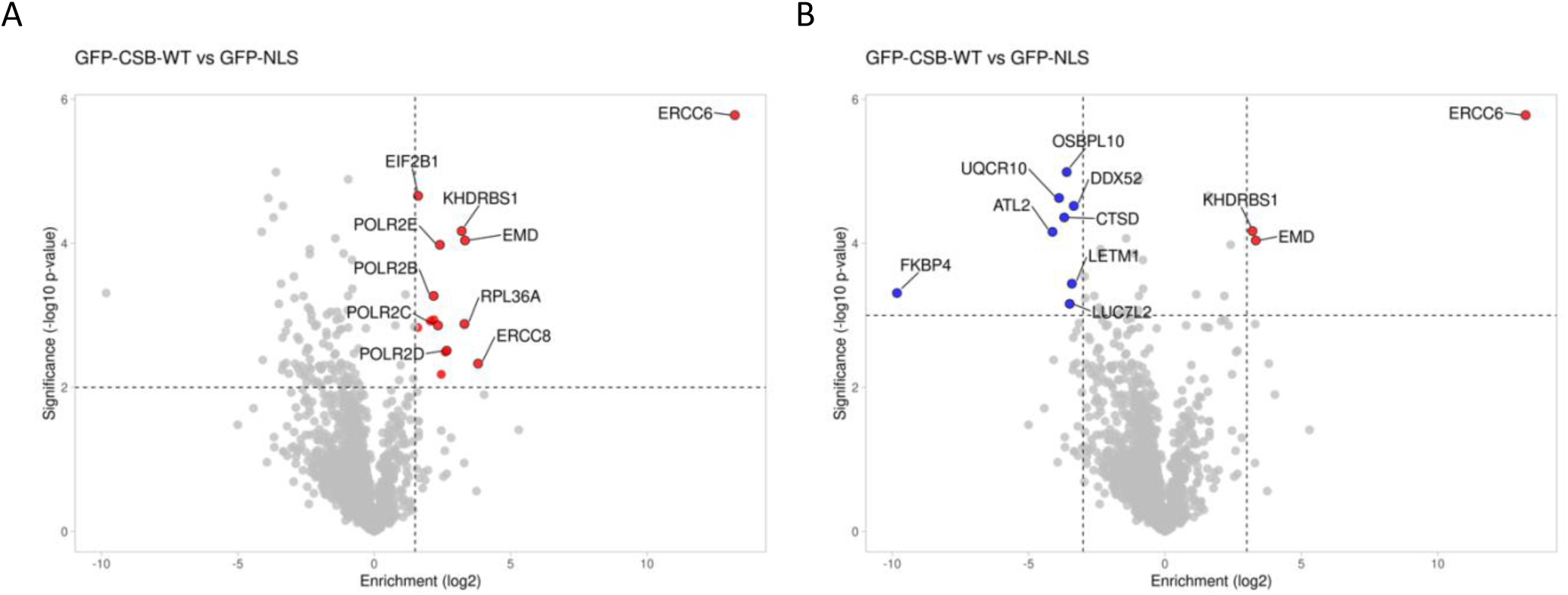
(A) Volcano plot generated from a published paper (van der Weegen et al., 2020). The interactive plot and all of the data is accessible through <u>this hyperlink</u>. (B) Volcano plot from the same data as (A) but replotted to show both significantly increased and decreased hits with the Enrichment and Significance threshold set to 3 and with a different color scale. The plot is accessible through <u>this hyperlink</u>.

Users can easily access the data and plot through this URL, inspect the data and replot it. Suppose that a user is interested in both showing and annotating increased and decreased proteins with more stringent threshold levels, the user can replot the data as shown in figure 2B. The URL can be copied and shared. This URL would be:

https://huygens.science.uva.nl/VolcaNoseR/?data=5;;Difference_CSB_GFP;p_value_CSB_GFP;Gene_names&vis=4;0.8;-3,3;3;significant;manh&can=20;;;&layout=;;;;;X;600;800&color=3;none&label=TRUE;GFP-CSB-WT%20vs%20GFP-NLS;TRUE;Enrichment%20(log2);Significance%20(-log10%20p-value);;24;24;18;6;&url=https://zenodo.org/record/3713174/files/CSV_1-GFP-CSB-WT_vs_GFP-NLS.csv

A list of settings that can be stored in the URL is available in a supplemental document (S1 text).

### Data re-use

To demonstrate the re-use of data, we examined the results of a recently published genome-wide CRISPR-based proliferation screen in a retina pigment epithelial (RPE1) cell line (Drainas et al., 2020). First, we retrieved the data of the 2D proliferation screens in wildtype and TP53 knockout cell lines (shown in figure 1B of the paper). The data of each of the screens is converted to CSV file and deposited at zenodo.org, doi: <u>10.5281/zenodo.3843685</u>. Next, we used the CSV file as input for VolcaNoseR and inspected the volcano plot (Figure 3). Given our interest in G protein-coupled receptor signaling (Chavez-Abiega et al., 2019), we looked for components of this signaling module. The GNAS gene was among the significant hits in the 2D proliferation screens in both wildtype and TP53 knockout cells (figure 3A and 3B), suggesting that it has an antiproliferative role in RPE1 cells. This result is in line with recent work on GNAS in the context of sonic hedgehog signaling (Pusapati et al., 2018b; a). This finding nicely demonstrates that the re-use of data from genome-wide, CRSIPR-based screens is an efficient way to generate or confirm hypotheses. Here, we show that the VolcaNoseR web tool can be used to mine current datasets, communicate new observations (figure 3), which can be easily shared through hyperlinks for re-use.

**Figure 3:**
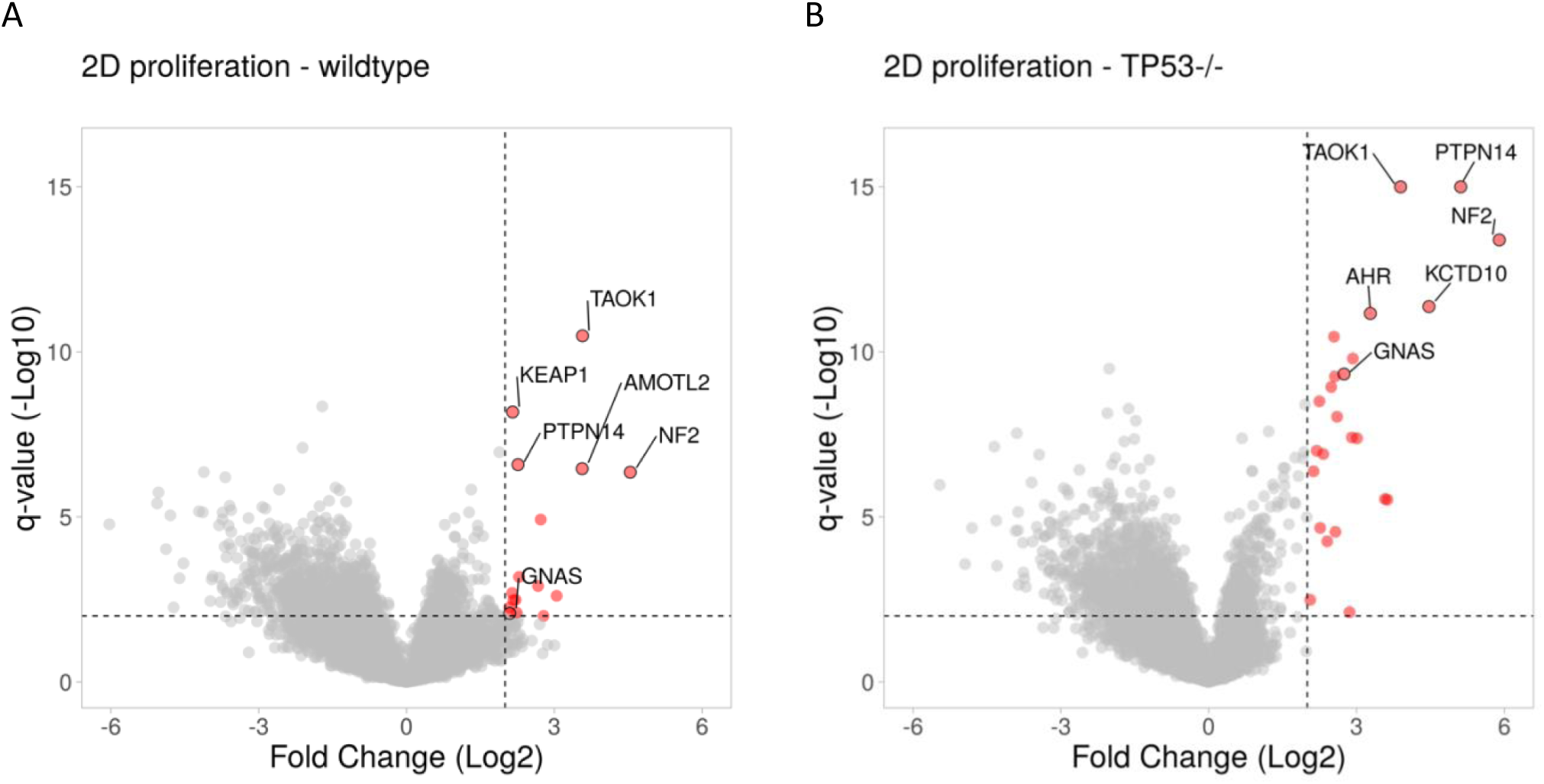
Volcano plots generated from a published dataset (Drainas et al., 2020) show that GNAS has an antiproliferative role in RPE cell growth. The volcano plots were generated with VolcaNoseR using published data and setting the threshold for the Log2(Fold Change) and -Log10(q-value) to 2. Significant hits are depicted in red and reflect genes that have an antiproliferative function. The top five candidates and GNAS are labeled. (A) Results from a 2D proliferation assay on RPE cells, plot and data are accessible through <u>this hyperlink</u>. (B)Results from a 2D proliferation assay on RPE TP53-/- cells, plot and data are accessible through <u>this hyperlink</u>.

## Conclusion

Volcano plots are data visualizations that are based on large datasets. Unfortunately, only a fraction of the data is labeled in static figures and, therefore, the vast majority of the information is inaccessible. To provide access to all of the data represented in a Volcano plot, we developed an interactive online plotting tool. By hovering over the plot with a pointer, each data point can be inspected. In addition, user-defined genes or proteins can be labeled in the plot and listed in a table. Together, these features enable access to all the information that the plot is based on. Finally, the web app can be used to share the data and the plot to allow other users to interact with the data and reuse it. Therefore, VolcaNoseR increases the transparency and re-use of large comparative genome- and proteome-wide datasets.

## Acknowledgments

Some of the VolcaNoseR code is taken from PlotTwist and it is partially inspired by the VolcanoR app (https://github.com/vovalive/volcanoR). We are grateful to Graham Dellaire (Dalhousie University, Canada) and Inés Pineda-Torra (University College of London, UK) for their input and thank Auke Folkerts (UvA, The Netherlands) for help with the server that runs shiny. The feedback, suggestions, enthusiastic responses and example plots that are shared on Twitter are highly appreciated.

