## Supplemental text S1 for "VolcaNoseR – a web app for creating, exploring, labeling and sharing volcano plots"

**S1 Text – Passing parameters to VolcaNoseR through the HTML address**

Passing parameters through the HTML address can be used to define a visualization and alter the standard layout. There are several queries (?data, ?vis, ..) that can be used. Each of the queries can hold multiple parameters. The queries are separated by an ampersand (&) and the parameters are separated by a semicolon(;).

Parameters that need no change (i.e. remain default) should be left empty. The parameters are defined by their position so their order is critical. Below, the parameters than can be changed through the HTML address are indicated, with the options between brackets:

?data

[1] data_input	{1/2/3/4/5}

[2] tidyInput	{T}

[3] x_var	variable (text)

[4] y_var	variable (text)

[5] g_var	variable (text)

?vis

[1] pointSize	{0-10}

[2] alphaInput	{0-1}

[3] fc_cutoff	Nummeric input from double slider, e.g.: -2,2

[4] p_cutoff	Nummeric input from slider, e.g.: 2

[5] direction	{all | significant | increased | decreased}

[6] criterion	{manh | euclid | fc | sig

?can

[1] top_x	{numeric}

[2] show_table	{T | F}

[3] hide_labels	{T | F}

[4] user_gene_list	list of comma-separated names

?layout

[1] rotate_plot	{T | F}

[3] change_scale	{T}

[4] range_x	-> number (minimum value),number (maximum value)

[5] range_y	-> number (minimum value),number (maximum value)

[6] For compatibility	‘X’

[7] plot_height	-> number reflecting pixels

[8] plot_width	-> number reflecting pixels

?color

[1] adjustcolors	{1 | 3 | 5}

[2] user_color_list	-> list of colors separated by comma’s

?label

[1] add_title	{T}

[2] title	-> text

[3] label_axes	{T}

[4] lab_x	-> text

[5] lab_y	-> text

[6] adj_fnt_sz	{T}

[7] fnt_sz_title	-> number

[8] fnt_sz_labs	-> number

[9] fnt_sz_ax	-> number

[10] fnt_sz_cand

[11] add_legend

?url

query[[‘url’]] URL	-> URL
